# Decoupling between causal understanding and awareness during learning and inference

**DOI:** 10.1101/391938

**Authors:** Jihyea Lee, Jerald D. Kralik, YuJin Cha, Suyeon Heo, Sang Wan Lee

**Affiliations:** Department of Bio and Brain Engineering, Korea Advanced Institute of Science Technology (KAIST) Daejeon 34141, Republic of Korea; Brain and Cognitive Engineering Program, Korea Advanced Institute of Science Technology (KAIST) Daejeon 34141, Republic of Korea; KI for Health Science Technology, Korea Advanced Institute of Science Technology (KAIST) Daejeon 34141, Republic of Korea; KI for Artificial Intelligence, Korea Advanced Institute of Science Technology (KAIST) Daejeon 34141, Republic of Korea

**Keywords:** inductive and deductive reasoning, causality, uncertainty, metacognition, cognitive psychology

## Abstract

Causal reasoning is a principal higher-cognitive ability of humans, however, much remains unknown, including (a) the *type* (systematic versus intermixed) and *order* (inductive-then-deductive or vice versa) of experience that best achieves causal-chain extraction; (b) how inferences generalize to novel problems, especially with one-shot experience; and (c) how metacognition, reflected in *uncertainty* of one’s knowledge, relates to actual knowledge. We tested people on a realistic cancer biology task (e.g., ‘seroc’ chemicals inducing tumors with subsequent effects). Systematic experience was superior, with some evidence that the inductive-then-deductive order promoted stronger one-shot generalization. Notably, uncertainty was decoupled from actual knowledge, with the deductive-then-inductive group being *overconfident*, likely reflecting lack of awareness of the inductive component; while those with successful one-shot generalization held *lower confidence*, reflecting generalization with minimal experience, while remaining skeptical. Our findings clarify processes underlying causal reasoning, and reveal a complex relationship between causal reasoning and metacognitive awareness of it.

## Introduction

Learning is the act of acquiring new or modifying existing knowledge, which frequently requires learning a relationship between different factors (e.g., stimuli, behaviors, outcomes). Although observations of relationships are inherently associational, *causality* may be captured formally by considering the degree of certainty one has that one factor will lead to another (Lee et al. 2015): when one is reasonably certain, it appears causal. Two distinct learning strategies for identifying relationships are incremental and one-shot learning^1,2,3^. Acquisition of knowledge that occurs gradually through repeated trial and error has been referred to as *incremental learning*; whereas, the phenomenon in which animals rapidly learn from only a single pairing of two factors such as a stimulus and consequence is called *one-shot learning*. A classic example for the latter is conditioned taste aversion: i.e., when the ingestion of a novel substance (conditioned stimulus, CS) is paired with internal malaise (unconditioned stimulus, US), an association between the taste and sickness can be quickly established, even with just one pairing, and even with long delays (4-12 hours) between them^4,5,6,7,8,9,10^.

Although taste aversion may be a specialized case^11,12^, it is clear that a substantial amount of human learning occurs via one-shot: i.e., after just one observation of the phenomenon. In fact, for one-shot learning we can distinguish two types: the learning of an entirely new problem (such as a novel stimulus-outcome pairing as in taste aversion), and the other via generalizing from prior knowledge based on similarity, such as one-shot categorization, in which people quickly learn to recognize novel objects using prior knowledge of object categories^13^.

Progress has also been made in determining when each learning strategy is employed. For example, there is evidence that both learning strategies are driven by prediction error, with higher learning rates reflecting faster and ultimately one-shot learning, and lower rates, the slower incremental strategy^1,2,14,15,16,17,18^. Moreover, evidence also supports the existence of a separable control process that sets the learning rates (of all possible relationships to be learned), and guides further processing in the brain that underlies the given learning strategy^1,19^. Finally, evidence suggests that the learning rate for any given case (such as a given stimulus-outcome pairing) is set by the control process based on the uncertainty of the relationship, with higher rates assigned to less certain pairings — that is, if a particular relationship is unclear, put extra effort into resolving it more quickly^1^.

Thus, in general, when uncertainty of a given relationship is high, one-shot learning is more likely to be deployed by the control process, with increased learning rates sufficient to resolve the uncertainty and learn the relationship. However, when uncertainty is high, an additional challenge may exist: the posed problem could be too difficult to solve. What then should the control process do if simply increasing the learning rate does not suffice? In fact, we can look to the standard scientific process for clues as to what, in principle, should be done. Causal factors (i.e., independent variables) must be isolated from confounds for each phenomenon (i.e., dependent variable) of interest, and achieving this generally requires *systematic experience* with the relationship in question (leading to actual experimental manipulation when possible)^20,21^. One might then assume that causal reasoning may also benefit from systematic experience, especially in more challenging settings.

We therefore attempted to extend the previous research to a more difficult real-world problem that also reflects everyday scenarios for people. In most real-world cases, causal relationships exist in sequences or chains — some *x* leading to some *y*, and in turn *y* leading to *z*, and so forth — and to comprehend these causal chains, two main types of reasoning are typically needed: *inductive* and *deductive inference* ^22,23,24,25^. For inductive, it involves identifying the causal factor among multiple possibilities (e.g., *x* causing *y* vs. possible *a*, *b*, *c* and others as causes); for deductive, comprehending a series of relations (*x* leading to *y*, *y* leading to *z*) as a sequence, with the transitive understanding that *x* would lead to *z*. Two critical and open questions, then, are whether a particular *type* and *order* of experience with the inductive vs. deductive components would facilitate comprehension of the entire causal chain: i.e., whether systematic or intermixed, and whether inductive then deductive or vice versa. In the current study, we therefore tested the effects of experience type and order. We did so on both causal learning and inference, and under both incremental and one-shot experiences.

Indeed, one of the important advantages of logical reasoning is applying the acquired knowledge to otherwise novel instances; but the mechanisms underlying this process remain underspecified^19,24^. In one case that focused on transitive inference, people were first trained on seven pictures of galaxies that formed a ranking based on age (i.e., older to younger: A > B > C, etc.)^24^. Their ability to transfer this ordered ‘schema’ to novel cases was then tested by exposing the participants to novel galaxies, training them with adjacent pairings of the novel galaxies with prior ones (e.g., A > H, H > B), then testing their knowledge with nonadjacent pairings. They found that those with a stronger grasp of the original hierarchy transferred most successfully. In the current study, we attempted to extend this work in multiple ways, including testing causal understanding beyond simple rank, both deductive and inductive components, and transfer in the extreme one-shot case: i.e., one trial of one novel exemplar.

At the same time, heightened problem difficulty poses additional challenges for the control system. Evidence for a control process that uses uncertainty to toggle among different learning strategies may suggest that uncertainty should closely match the actual likelihood of problem-solving accuracy. However, exactly how the uncertainty assessment achieved is not fully known, especially in cases with heightened problem difficulty. Since the uncertainty assessment is based on a separable process, it could in principle have a more complex relationship with actual problem-solving accuracy, with possible mismatches including overconfidence (knowing less than you think) or lack of it (knowing more than you think). And yet such possible decoupling may be difficult to produce with less challenging problem paradigms. The final major aim of the current study, therefore, was to examine this relationship more closely.

In sum, we designed the current study to test (a) the *type* (sequential vs. intermixed) and *order* (inductive then deductive vs. vice versa) of experience that best achieves extraction of causal chains requiring inductive and deductive reasoning; (b) how inferences generalize to novel problems from single *one-shot* experience with the novel case; and (c) how metacognition, reflected in *uncertainty* of one’s knowledge, relates to actual problem knowledge. We tested people on a cancer biology task (e.g., ‘seroc’ based chemicals inducing tumors with particular effects) designed to provide a realistic causal reasoning problem.

## Results

We constructed sentence stimuli that, based on inductive and deductive inference, formed causal chains of events (Table 1, Fig. 1). For example, using inductive inference, if substances that induce the tumor ‘Karmeictumor’ include Acoseroc and Benzoseroc, one could infer that all substances containing ‘–seroc’ in their name are likely causal. And using deductive inference, if ‘Acoseroc induces the tumor Karmeictumor’, ‘Karmeictumor triggers neckache’, and ‘tumors that trigger neckache eventually metastasize’ then ‘Acoseroc can lead to tumor metastasis’. Construction of the complete causal chain (Fig. 1) was referred to as *extracting the rule*. We then asked whether the *type* (sequential vs. intermixed) and *order* (inductive then deductive vs. vice versa) of experience with the sentences affected learning and inference, both after an initial incremental learning condition and after a subsequent one-shot experience with a novel causal chain. In addition, we examined the relationship of actual causal knowledge to the participants’ metacognitive assessment of their knowledge via *confidence* in their answers.

**Table 1.**
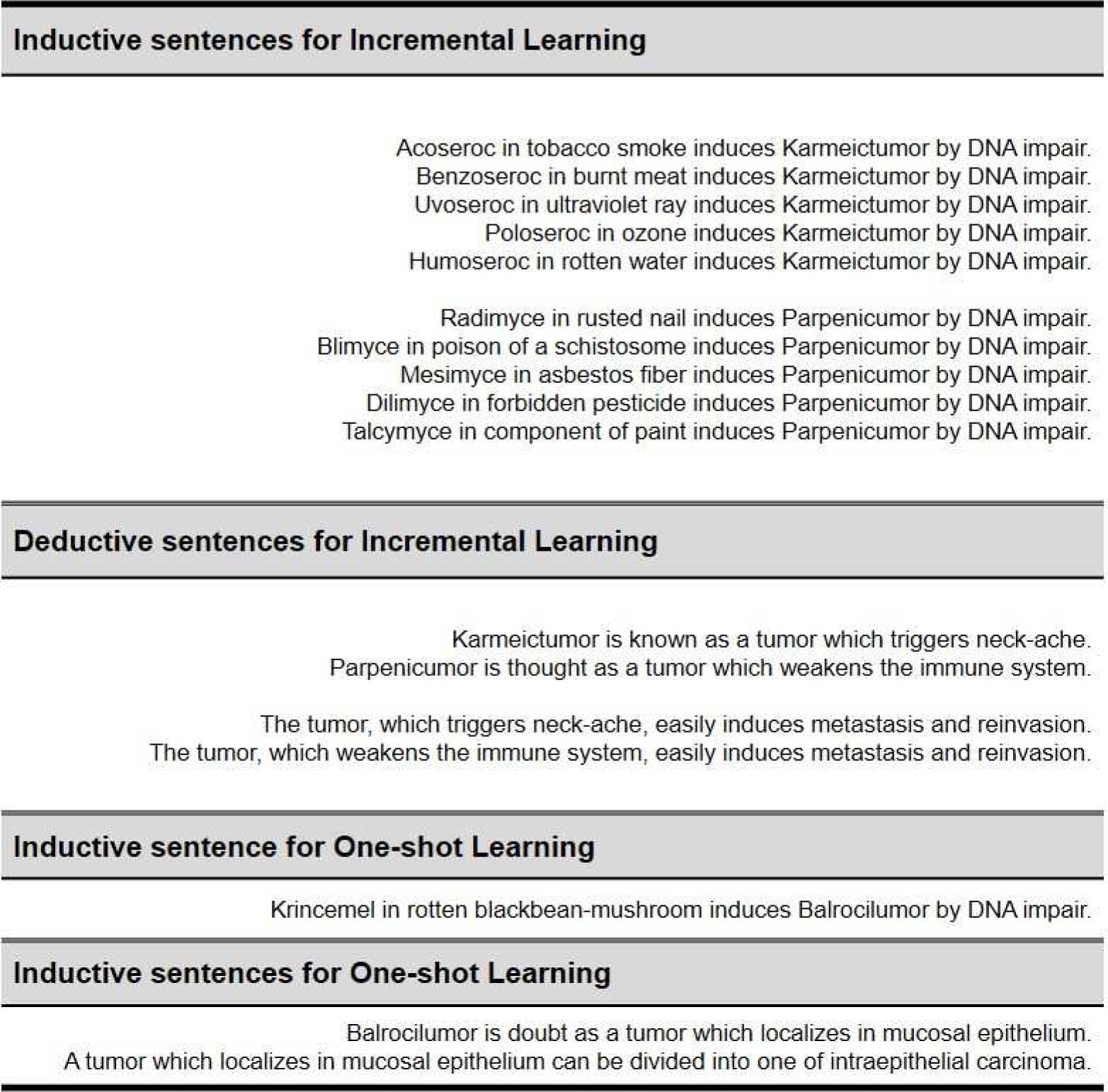
Sentence stimuli. All of the stimuli were presented in Korean. See the ‘Task Paradigm’ section for a description of how the inductive sentences for incremental learning also included a “did not induce” structure, used to increase difficulty and uncertainty.

**Figure 1.**
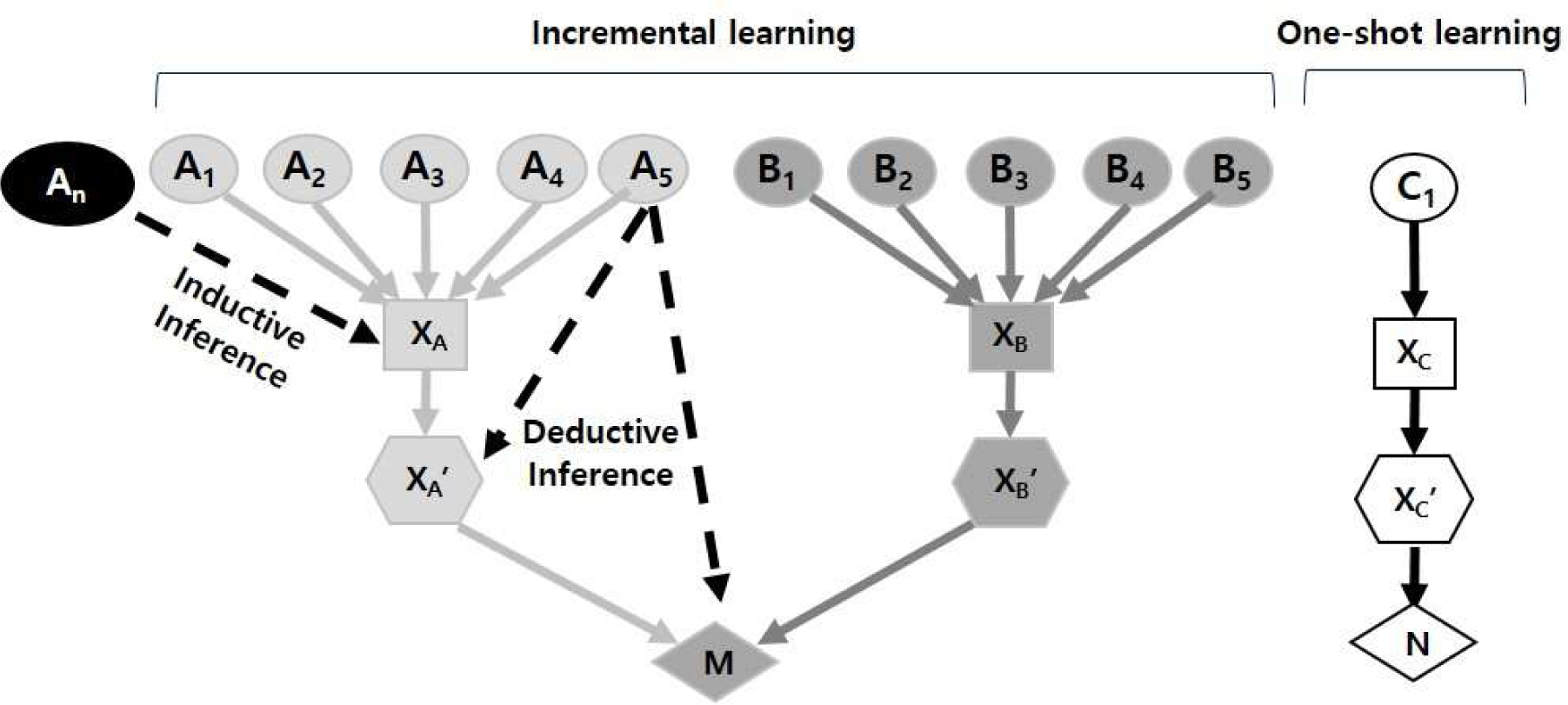
The sentence structure used for test stimuli. For inductive sentences for Incremental learning, A_1_ – A_5_ and B_1_ – B_5_ represent the specific substances that induced tumor growth (e.g., A_1_ = Acoseroc, B_1_ = Radimyce); while X_A_ and X_B_ represent the tumors (Karmeictumor, Parpenicumor). For deductive sentences for Incremental learning, X_A_’ and X_B_’ represent neck-ache and weakened immune system respectively, while M represents the final outcome (metastasis and reinvasion). A_n_ represents the inductive inference to be drawn from A_1_ – A_5_ (suffix ‘-seroc’); B_n_, not depicted, is the suffix ‘-myce’. The dashed lines from A_5_ to X_A_’ and M reflect the deductive inferences that can be drawn, with those for A_1_ – A_4_, B_1_ – B_5_, A_n_, and B_n_ not depicted. The structure for one-shot learning was the same, with C_1_ representing both the specific substance used (Krincemel) and the actual causal suffix ‘-cemel’, X_C_ representing Barlocilumor, X_C_’ representing mucosal epithelium, and N representing intraepithelial carcinoma. See Table 1 for specific sentences.

To address these research questions, we divided the participants into three groups: (1) those that received inductive-then-deductive incremental experience, (2) vice versa, and (3) those who experienced both the inductive and deductive components simultaneously — i.e., randomly across trials. All three groups then experienced the one-shot trial with the novel problem (with inductive and deductive components in the same order as in the incremental condition—see Materials and Methods). The participants then received a 35-question test to assess their knowledge of (a) the specific sentences presented (called “Learning” questions), and (b) the possible inferences made (called “Inference” questions). Thus, the test questions fell into eight categories: Incremental Inductive Learning, Incremental Inductive Inference, Incremental Deductive Learning, Incremental Deductive Inference, One-shot Inductive Learning, One-shot Inductive Inference, One-shot Deductive Inference, and One-shot Inductive + Deductive Inference. See Materials and Methods for details.

Finally, to examine the relationship between actual causal reasoning (i.e., performance on the test questions) and one’s own assessment of their knowledge, for all test questions we also asked the participants to rate the certainty of each answer from −5 to 5.

### Incremental inductive learning and inference leads to overconfidence

#### Incremental learning and inference

Overall, the total mean scores for all questions combined revealed no significant differences among the three groups; however, the difference in confidence among the groups was significant (ANOVA, F=4.304, p<0.05), with mean confidence of Group 2 being significantly higher than that of Group 3 (p<0.01) (Fig. 2). Thus, deductive-then-inductive experience led to greater confidence in test answers over random order, although the heightened confidence was not reflected in actual performance, at least overall.

**Figure 2.**
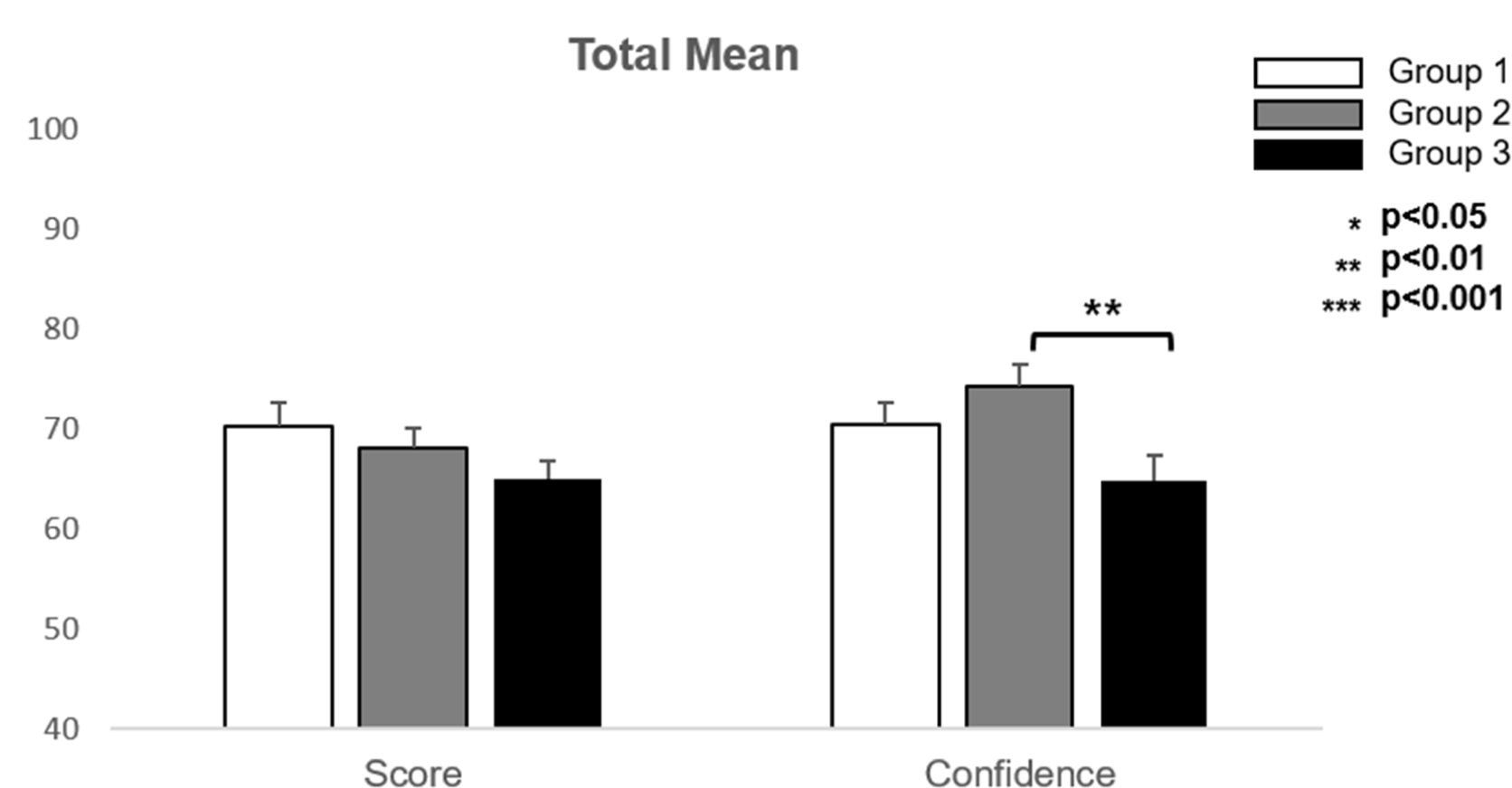
Total mean score and confidence of each group. The mean confidences values were significantly different among all three groups (p<0.05). The mean confidence of Group 2 (mean 74.336, SE 2.11) was higher than that of Group 3 (mean 64.738, SE 2.61); p<0.01.

To examine these findings more closely, we next analyzed the results for each of the main question types, focusing first on incremental learning and inference (Fig. 3). Examining performance scores first, and focusing first on learning, in which questions were based on the actual sentences used (Table 1), we found no overall difference among the groups in the inductive learning performance scores (Fig. 3A), but we did find a difference for deductive learning (ANOVA, F=3.991, p<0.05), with Group 1 having a significantly higher score than Group 2 (p<0.01; Fig. 3C), indicating that receiving the deductive training last promoted retention of the specific training exemplars used for deductive learning.

**Figure 3.**
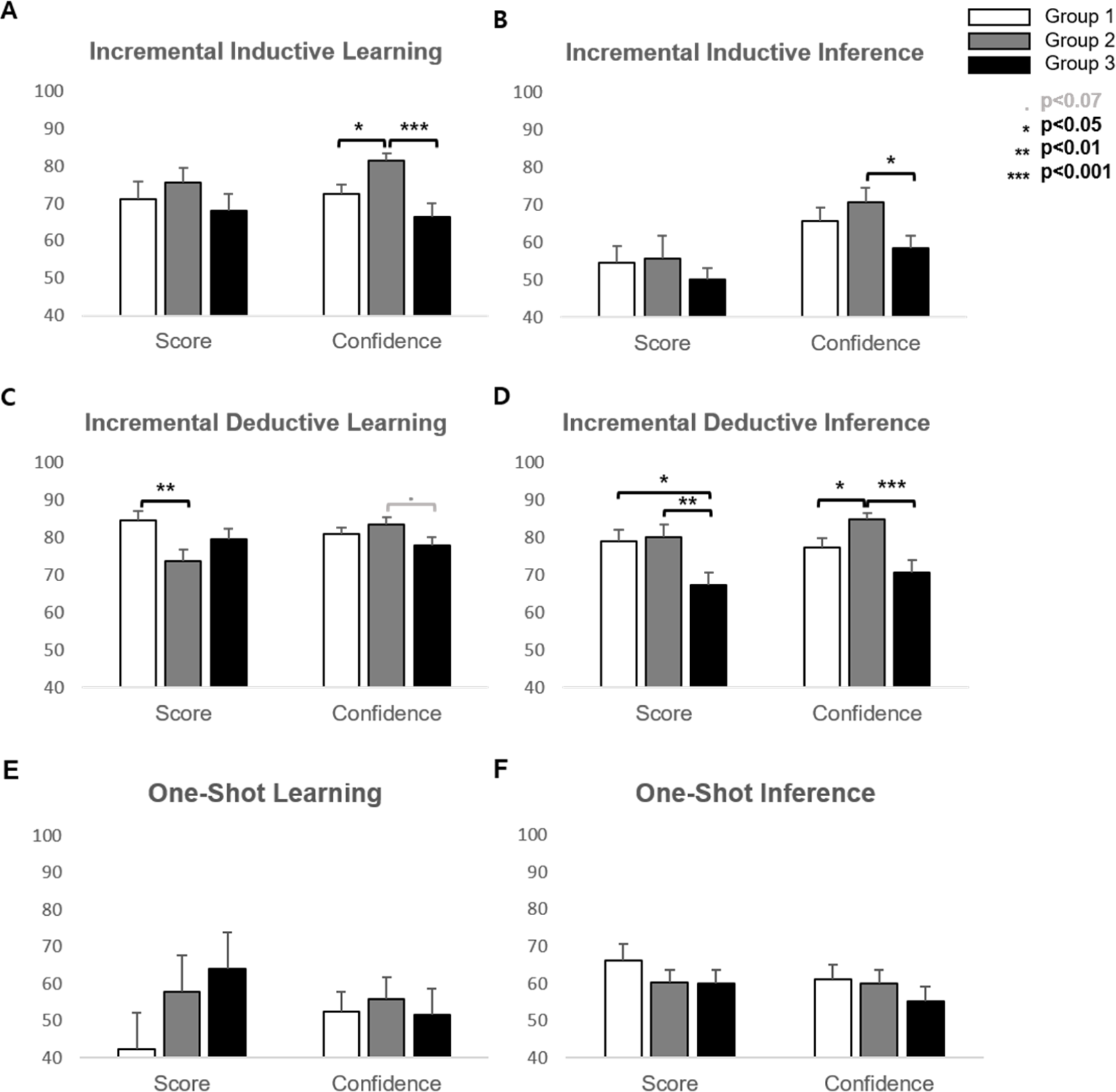
Performance scores and confidence ratings of each group when the questions divided into several types. **(A)** The confidences of IC Inductive Learning questions showed difference between all three groups (p< 0.01). Group 2 (mean 81.410, SE 2.07) had higher confidence than both Group 1 (mean 72.435, SE 2.70; p<0.05) and Group 3 (mean 66.303, SE 3.65; p<0.001). **(B)** In the case of IC Inductive Inference questions, confidence of Group 2 (mean 70.746, SE 3.74) recorded higher than Group 3 (mean 58.424, SE 3.34) with p<0.05. **(C)** shows the case of IC Deductive Learning that score differences between all three groups were significant (p<0.05). Score of Group 1 (mean 84.615, SE 2.37) was higher than that of Group 2 (mean 73.626, SE 3.04) with p<0.01. Difference of confidences between Group 2 (mean 83.467, SE 1.95) and Group 3 (mean 77.766, SE 2.29) was close to being statistically significant (p-value=0.06). **(D)** For IC Deductive Inference part, score and confidence both showed significant differences between all three groups with p<0.05, p<0.001 each. Scores of Group 1 (mean 78.846, SE 3.14) and Group 2 (mean 80.128, SE 3.21) were higher than score of Group 3 (mean 67.333, SE 3.26) with p<0.05, p<0.01 respectively. In confidence, Group 2 (mean 84.732, SE 1.74) got more confident than Group 1 (mean 77.389, SE 2.43; p<0.05) and Group 3 (mean 70.545, SE 3.32; p<0.001) each. **(E), (F)** show the case of OS Learning and Inference that there were no significant differences between groups.

For incremental inductive inference, we also found no significant differences among the groups on the incremental inference scores (Fig. 3B), but we did find differences for deductive inference, with Groups 1 and 2 both obtaining higher scores than Group 3 (p<0.05, p<0.01; Fig. 3D). With scores near 50% (and thus near chance) for inductive inference, it is clear that the inductive inference test was difficult for the participants; while the deductive inference component was easier, at least for Groups 1 and 2 (Fig. 3D).

In contrast, and as reflected in the overall results (Fig. 2), the confidence ratings did not match the performance scores. Although we found no significant differences among performance scores for inductive learning (Fig. 3A), confidence was significantly higher for Group 2 than for both Groups 1 and 3 (p<0.05, p<0.001); and although Group 1 obtained a significantly higher performance score than Group 2 for deductive learning (Fig. 3C), with Group 2 actually having a lower score than Group 3, Group 2 nonetheless held confidence in their scores comparable to Group 1 and to a level greater than Group 3 that approached significance (p=0.06). For incremental inductive inference (Fig. 3B), in which all groups performed relatively poorly (indicating that the incremental inference problem component was particularly challenging), Groups 1 & 2 nonetheless exhibited relatively high confidence in their performance, with Group 2’s being significantly higher than that of Group 3 (p<0.05). For deductive inference (Fig. 3D), where the performance scores of both Groups 1 & 2 were relatively high, Group 2’s confidence nonetheless particularly stood out, exceeded that of both Groups 1 and 3 significantly (p<0.05, p<0.001). Thus, for incremental learning and inference, and for both the inductive and deductive components, we found a mismatch between performance scores and confidence in them, with the deductive-then-inductive experience order appearing to inflate confidence above actual performance. We consider this result in light of the entirety of the findings in the discussion.

#### One-shot learning and inference

For one-shot learning and inference, we found no differences among the groups for any of the question types, for either performance scores or confidence in performance — we thus show the results for One-Shot Learning (Fig. 3E) and One-Shot Inference (Fig. 3F) combined for the inductive and deductive components. In fact, the relatively low performance scores for one-shot learning and inference for all participants combined suggest that, overall, participants had difficulty with this task component, both with respect to remembering the specific information presented for the one-shot problem (Table 1), and transferring a rule from the incremental task component to the novel, one-shot problem exemplar.

### One-shot inference culminates in underconfident generalization

#### Incremental learning and inference

From the post-test interview, we could identify the participants who were able to infer the overall causal chain or *rule*. Because participants did well on the deductive component of the task, in the post-experiment interview we identified those who also successfully extracted the inductive component, and thus those who reported a general relationship between a suffix and a cancer disease state. To examine the results of these participants, we defined two new groups based on whether they inferred the rule, denoted Group O, and those who did not, denoted Group X. Four participants who extracted the rule but did not use it to solve the inference questions were excluded (one from Group 1, three from Group 2). The proportion of “rule-catch” participants in each group was 42.3%, 46.2% and 8%, respectively. This result has two main implications: first, for *type* of experience, prior experience with inductive and deductive learning and inference, regardless of order, promoted the ability to make the proper inferences and extract the causal chain, i.e., the rule; second, the *order* of inductive versus deductive learning did not significantly impact the number of participants who could infer the general rule.

Comparing the rule-catch and non-rule-catch groups overall, as expected Group O performed significantly better than Group X (p<0.01). However, although the confidence rating was higher for the group, confidence overall was not significantly different (Fig. 4). We next examined the results broken down by question type (Fig. 5). For the incremental experience condition, for both the inductive and deductive *learning* questions, i.e., based on the specific exemplars experienced (Table 1), there were no differences between Groups X and O for either performance score or confidence (Fig. 5A&C). These findings reflect the fact that the division between groups was based on their inference success, and further suggest a relative independence of the learning and inferencing components of the problem, given the different effects. For both inductive (Fig. 5B) and deductive (Fig. 5D) inference, in contrast, we found significant differences between the groups for both performance scores and confidence (In inductive inference; score p<0.001, confidence p<0.05 and in deductive inference; score p<0.05, confidence p<0.05). Thus, in this case, the performance scores and confidence ratings of the participants were aligned, reflecting actual problem understanding: i.e., the ability to infer the general rule from the original exemplar sentences.

**Figure 4.**
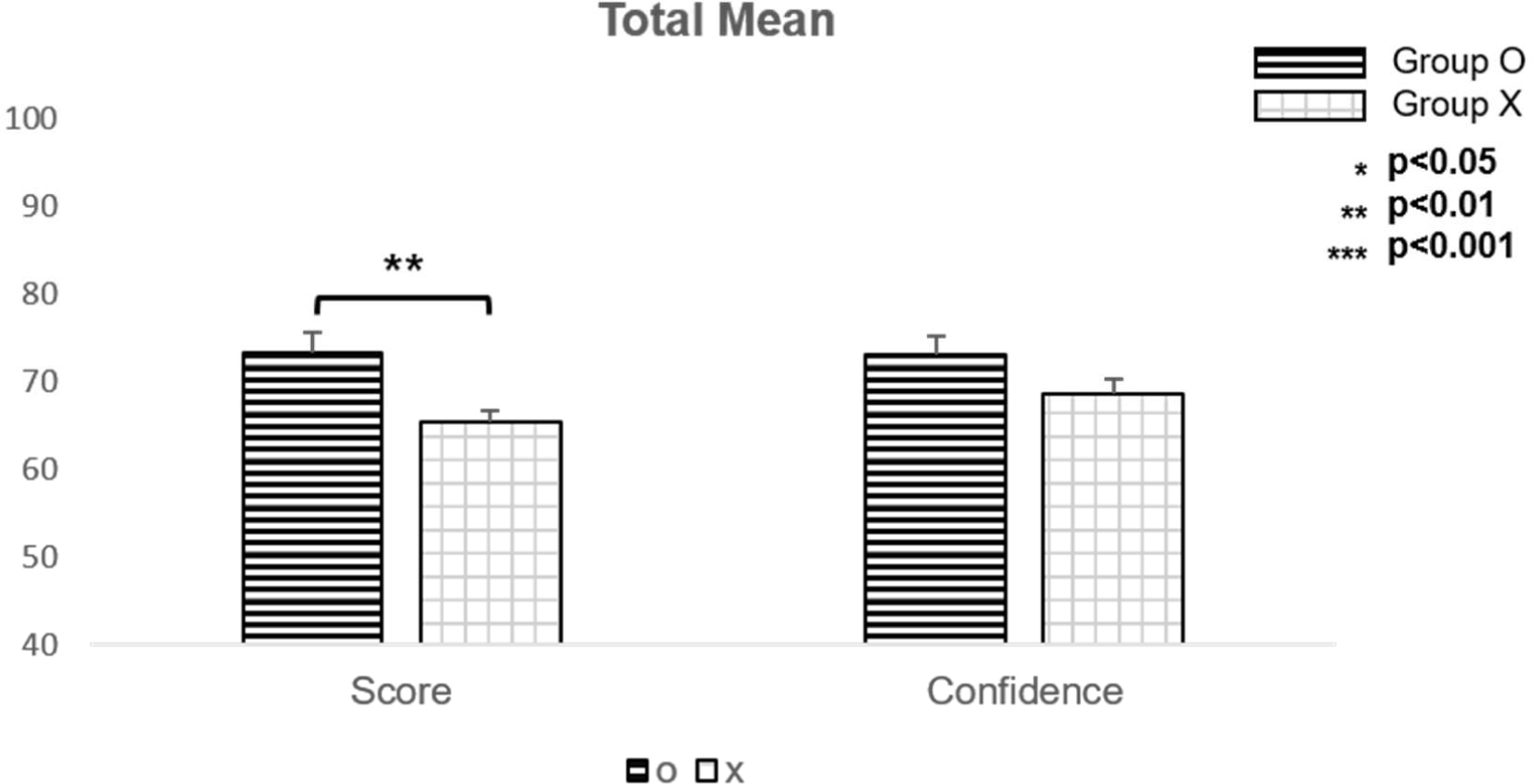
Total mean score and confidence of each group divided by rule-catch. The mean score of Group O (mean 73.214, SE 2.25) was significantly higher than that of Group X (mean 65.337, SE 1.35); p-value<0.01.

**Figure 5.**
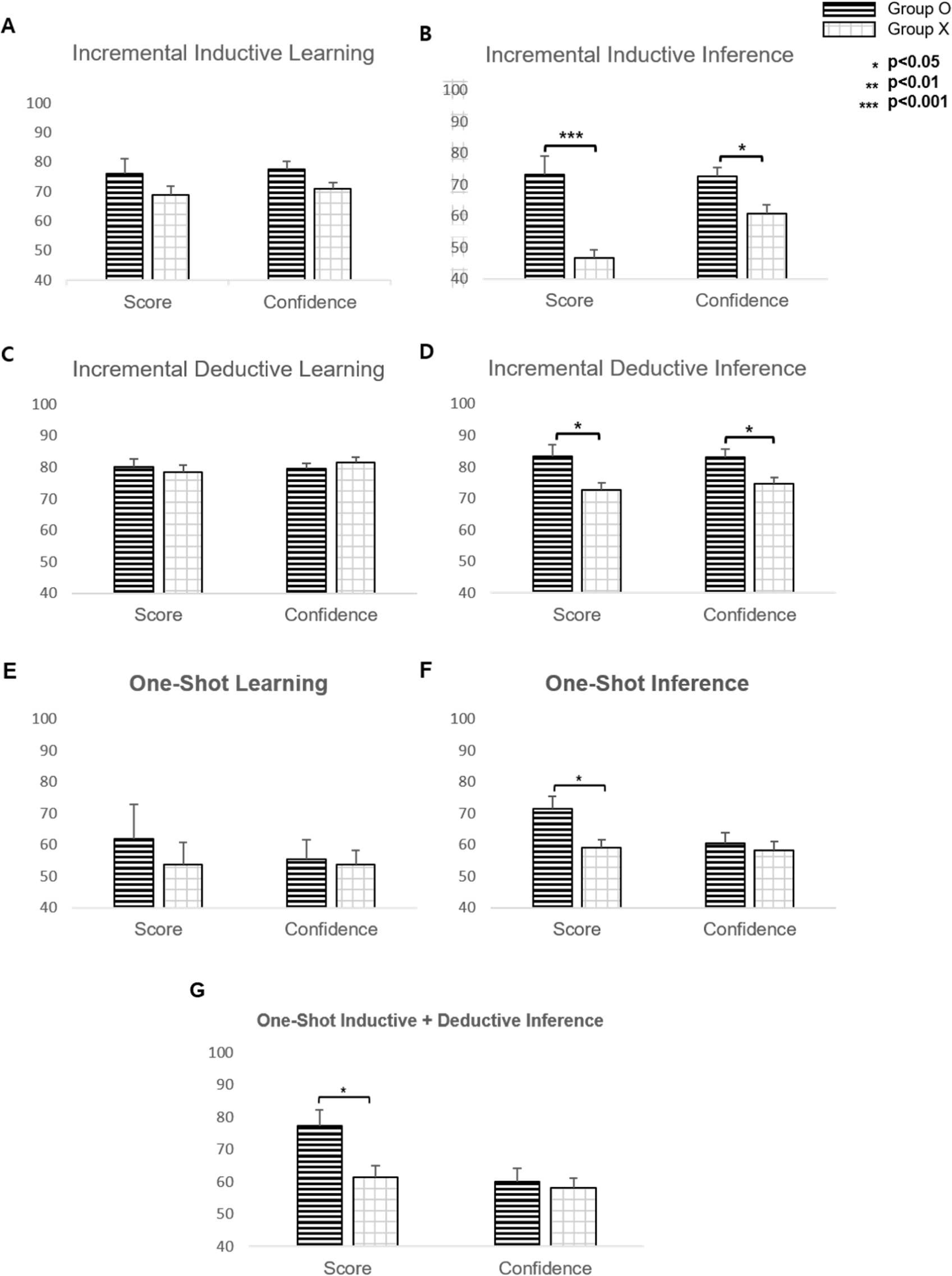
Performance scores and confidence ratings of two groups based on rule-catch when the questions divided into several types. **(A)** The difference in scores and confidences of IC Inductive Learning part between two groups were not significant. **(B)** In the case of IC Inductive Inference questions, both score and confidence of Group O (score mean 73.016, SE 6.03; confidence mean 72.583, SE 2.86) recorded higher than Group X (score mean 46.795, SE 2.47; confidence mean 60.897, SE 2.63) with p<0.001 and p<0.05 respectively. **(C)** The result of IC Deductive Learning shows no significant differences. **(D)** For IC Deductive Inference part, score and confidence both showed significant differences between two groups with p<0.05 both (Group O score mean 83.333 SE 3,64; confidence mean 83.117, SE 2.46; Group X score mean 72.756, SE 2.24; confidence mean 74.738, SE 2.02). **(E)** In OS learning, differences were not significant. However, **(F)** In OS Inference, Group O had a higher score (mean 71.429, SE 4.03) than Group X (mean 58.974, SE 2.70); p<0.05. **(G)** Performance scores and confidence ratings of two groups based on rule-catch with the questions about OS Inductive + Deductive Inference. Group O (mean 77.381, SE 4.85) was higher than that of Group X (mean 61.538, SE 3.46); p-value<0.05.

#### One-shot learning and inference

For one-shot *learning*, we again did not find a significant difference between the two groups either for performance score or confidence (for either inductive or deductive components) (Fig. 5E). However, for both one-shot inference (i.e., inductive and deductive inference questions combined) (Fig. 5F) and one-shot inductive + deductive inference (i.e., questions probing understanding of the entire causal chain) (Fig. 5G), the performance score for Group O was significantly higher than that of Group X (p<0.05). The results thus indicate that actually abstracting the inference rule was critical for solving the inference problem we constructed, and classifying participants based on the follow-up interview accurately reflected this successful abstraction. Moreover, additional evidence for this successful abstraction process comes from the significant difference between the O and X groups with the *inference* as opposed to *learning* questions (Fig. 5E vs. 5F & G). Yet at the same time, the successful abstraction of the rule and relatively high performance scores did not lead to significantly higher confidence in their one-shot performance, with confidence ratings remaining comparable to the non-rule-catch group. Thus, even though the Group X participants performed relatively well by transferring the extraction of the rule from the incremental experience to the one-shot exemplar, their confidence in this transfer remained low: i.e., transfer occurred, but they appeared nonetheless skeptical about it. Thus, as opposed to the incremental learning and inference case, in which those experiencing the deductive-then-inductive order held overinflated confidence compared to their actual performance scores, and as opposed to the rule-catch group’s confidence with incremental inference, which generally matched performance, in the one-shot experience case, the rule-catch group revealed a lack of confidence in their answers, even though they nonetheless generalized from their prior knowledge and made the proper inference.

#### Examination of experience order for rule-catch group only

Although the incremental sequential orders given to Group 1 (inductive-then-deductive) and 2 (vice versa) appeared to promote general rule inference comparably (42.3% and 46.2% rule-learners respectively), could the order nonetheless influence the strength of the successfully extracted rule and/or the ability to promote one-shot inference? To address these questions, we analyzed the performance scores, confidence, and one-shot inference ability of Group 1 versus 2 for only the rule-learners. We again found no evidence for order except in two cases, in which the Group 1 performance scores were higher than those of Group 2 for one-shot inference (i.e., inductive and deductive questions combined) (Group 1: 71.4 ± 4.0; Group 2: 59.0 ± 2.7) and one-shot inductive+deductive inference (i.e., questions that tested the entire one-shot causal chain) (Group 1: 77.4 ± 4.9; Group 2: 61.5 ± 3.5), with the difference in both cases approaching significance (stat 1, p<0.05; stat 2, p<0.05, respectively). Thus, we did find some evidence that receiving inductive experience prior to deductive may promote stronger rule extraction that leads to greater generalization of the inference rule to novel instances.

### Decoupling between performance and confidence during learning and inference

Finally, to provide a general, overall assessment of the interrelationships among the main factors of training order, test performance scores, confidence, and actual rule extraction, we used multiple linear regression. We conducted three regressions with test performance scores, confidence, and rule extraction as the dependent variable for each regression, respectively. For test scores and confidence ratings of them we used all test questions. For performance scores, the independent variables were ‘Group (x_1_)’, ‘Rule (x_2_)’ and ‘mean confidence (x_3_)’. For the Group variable in the regressions, because both sequential incremental learning orders (i.e., Groups 1 and 2) promoted the extraction of a general rule, we focused here on Groups 1 and 2 versus Group 3, so that Groups 1and 2 were coded as x_1_=1 and Group 3 coded as x_1_=2. The ‘Rule’ variable coded rule catching vs. not, with Group O as x_2_=1 and Group X as x_2_=2. A significant regression equation was found (F=5.738, p<0.01), with an adjusted R^2^ of 0.157. Participants’ predicted mean score was equal to 73.68 – 0.116x_1_ – 8.758x_2_ + 0.128X_3_. As shown in Table 2, ‘Rule’ proved to be the only significant predictor of mean performance score; in fact, when participants failed to catch the rule, the mean score decreased about 8.8. Thus, test performance was indeed a function of how well the participants inferred the general rule from the learning exemplars.

To determine the factors underlying confidence, we examined Group and Rule, as defined for the first regression above, as well as mean score set as the third independent variable x_3_. A significant regression was again found, with F=3.188, p<0.05, R^2^=0.079, and participants’ predicted mean confidence equal to 68.024 – 6.481x_1_ – 1.013x_2_ + 0.179x_3_. As also seen in Table 2, only the Group variable was found to be significant in this second regression, although the Group*Rule interaction was also significant. Thus, we again found that rather than confidence being driven largely by proper rule inference (Rule main effect) and thus actual understanding of the test material, it was driven more by prior history: i.e., receiving systematic incremental experience (i.e., the inductive and deductive components separately), rather than randomly experienced.

**Table 2.**
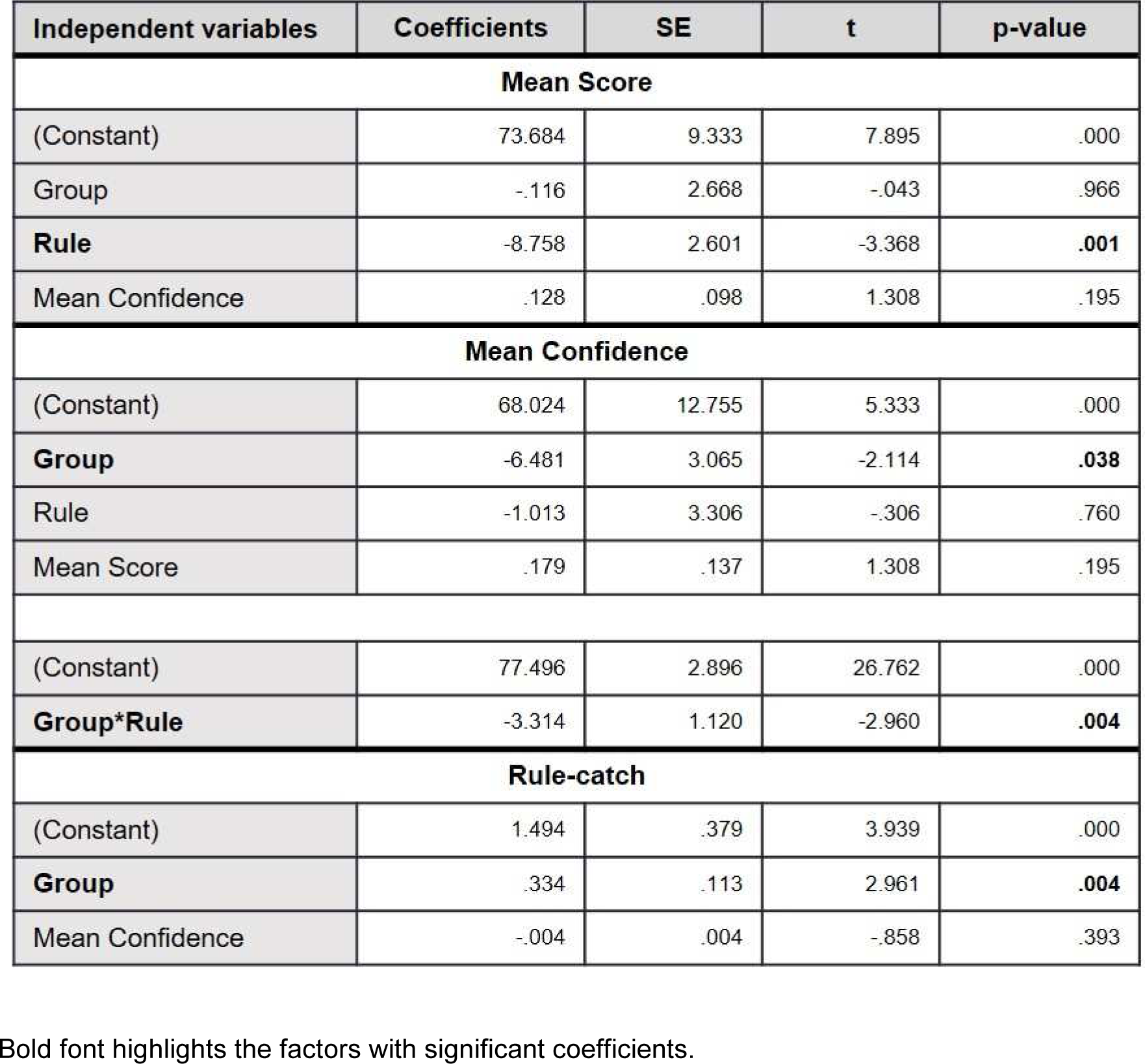
Results of multiple linear regression.

For the final regression, to determine what factors most enabled the participants to extract the general inference rule, we examined Group (x_1_) and mean confidence (x_2_) as independent variables. A significant regression equation was found with F=6.015, p<0.01, an adjusted R^2^=0.117. As seen in Table 2, the ‘Group’ variable was significant. Thus, as found throughout the study, the random presentation of the inductive and deductive components of the causal chains proved exceedingly difficult for participants to make further inferences from them; and thus participants needed to experience the inductive and deductive components separately to make the proper inferences.

Finally, Fig. 6 provides a summary diagram of the main regression findings: i.e., the relationship between Group, Rule, Score, and Confidence. In short, sequential incremental learning (i.e., Groups 1 and 2 vs. 3) promoted greater rule inference and greater confidence; when the rule was properly extracted, it was indeed reflected in the performance scores; and yet we did not find a direct, linear relationship between actual test scores and confidence in their test performance. In fact, we found evidence for a more complex relationship between them, with evidence for overconfidence, a proper level of confidence, and a lack of it, discussed further below.

**Figure 6.**
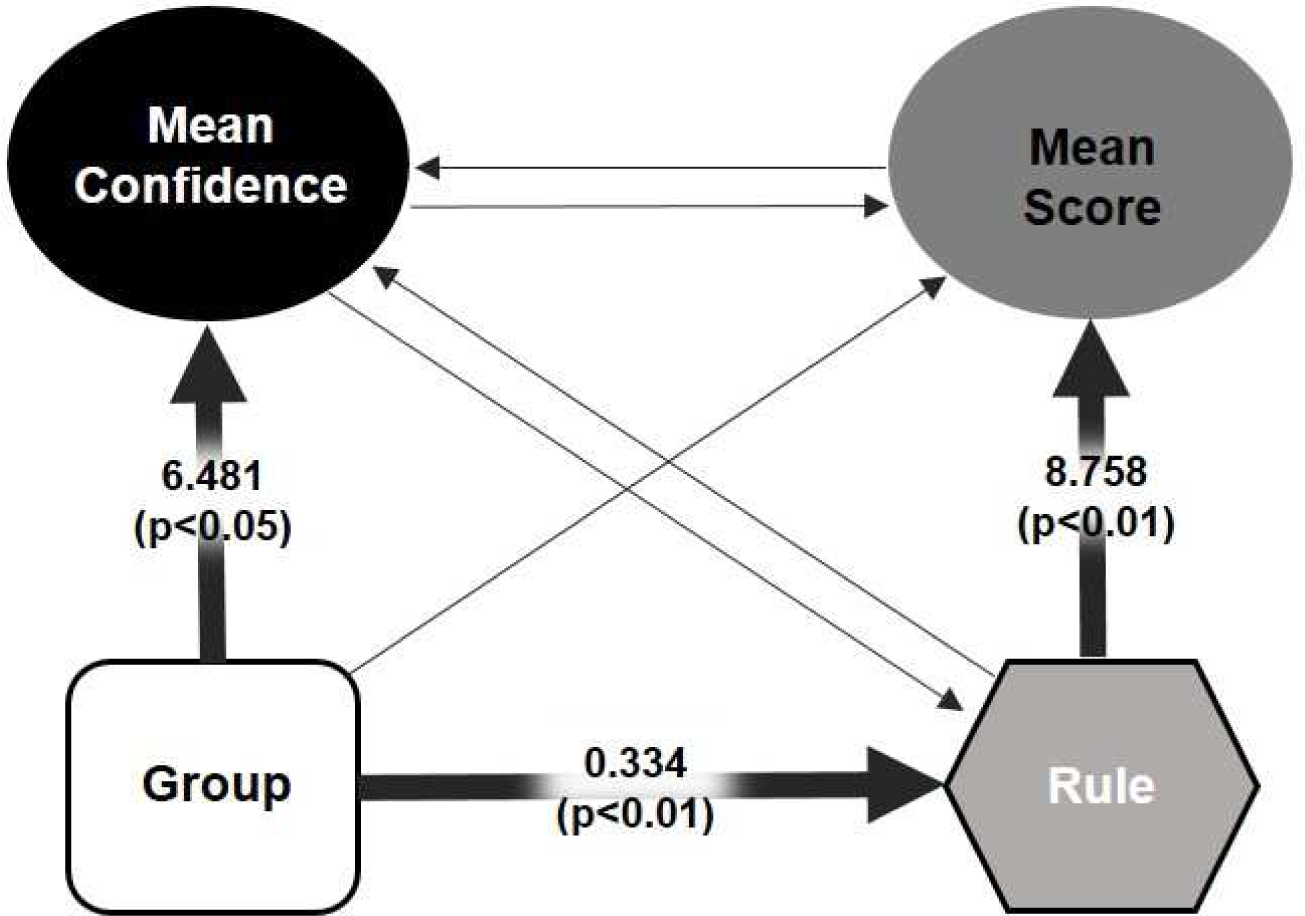
Relationships between experimental factors, performance and confidence level. The diagram was made based on the multiple linear regression results. Sequential incremental learning (i.e., Groups 1 and 2 vs. 3) promoted greater rule inference and greater confidence; when the rule was properly extracted, it was reflected in the performance scores; and yet we did not find a direct, linear relationship between actual test scores and confidence in their test performance.

## Discussion

Recent evidence suggests that different learning strategies may be employed based on uncertainty, with greater uncertainty (e.g., of how a given stimulus relates to an outcome) leading to higher learning rates, reflecting faster and ultimately one-shot learning, and lower rates reflecting the slower incremental strategy^1,15^. Thus, with relatively manageable problems, learning rate manipulation can be sufficient; but as problem difficulty increases, more elaborate means are likely needed for effective learning. In general, these may include more *systematic experience* with the factors to be learned and inferred, so that the problem becomes manageable, enabling successful extraction of the causal chains (i.e., logical inference). But what this ‘systematic experience’ may entail remains unclear. Here we focus on human high-level causal reasoning to determine whether the assembly of a complete causal chain via both inductive and deductive inference processes benefits by a particular type and order of experience (i.e., whether inductive-then-deductive, vice versa, or random). We also examined these effects on both incremental and one-shot experience, with the former providing multiple exemplars of a given rule with multiple trials of each, and the latter requiring the transfer of inference achieved from previous experience to a novel case consisting of one trial of one new exemplar. We tested these questions using realistic scenarios in cancer biology.

### Incremental experience

*For learning* (i.e., test questions based on the specific examples used in training), the results appeared to be highly driven by basic memory processes, revealing, for example, apparent recency effects in the incremental condition, in which the component experienced last (i.e., inductive or deductive) enjoyed more success.

For *inference*, the inductive component of the reasoning problem proved particularly challenging, and the only order effect we found among the specific reasoning types (inductive vs. deductive) was a detrimental effect of random order on deductive inference. Overall, however, it was clear that random order harmed inference, which was borne out in the third linear regression that focused on rule-catch (i.e., the factors enabling the ability to extract the rule properly, as identified by the post-experimental interview), with group (i.e., Groups 1 & 2 vs. 3) having a significant main effect on rule. Thus, isolated experience with the specific examples for each reasoning component (inductive and deductive inference) appeared necessary to see the problem components and their relationships clearly enough to enable proper inferences.

Yet for both learning and inference in the incremental experience condition, and for both inductive and deductive inference individually, the *confidence* of Group 2 (deductive-then-inductive experience) outpaced that for the other two groups, leading as well to a main effect of group on confidence both in the general ANOVA and in the second regression, even though no differences were found for the actual performance scores. Why would confidence, especially for the deductive-then-inductive order, be overinflated? Heightened confidence would be due to (a) an actual better understanding, leading to greater certainty, or (b) a false confidence, born from misunderstanding. Since the results revealed no significant differences in the level of understanding between groups, it suggests that Group 2 developed a false confidence.

Misunderstanding and subsequent false confidence could potentially reflect a lack of sufficient engagement in the problem to appreciate the difference between a general understanding and actual precise understanding of the details being tested. However, we assume that the participants were sufficiently motivated to engage sufficiently in the task details. A misunderstanding, then, may otherwise derive from focusing on wrong aspects of the problem, which nonetheless are confounded with the correct ones, resulting in a lack of negative feedback to signal misunderstanding. For example, when participants in Group 2 first received the deductive incremental learning component, they potentially learned the X_A_→X_A_’ and X_B_→X_B_’ links (see Figure 1), as well as X_A_’→M and X_B_’→M; and they presumably inferred the X_A_→M and X_B_→M links as well, with results showing that participants generally did well with the deductive component. Thus, they focused on various cause→effect links, and deductive inference. When they then moved to incremental *inductive* learning, it nonetheless included learning the deductive link at the beginning of the causal chain, i.e., A_1_→X_A_, for all A_1_-A_5_ and B_1_-B_5_. If participants learned the specific names of all 1-5, they would score well on the learning questions for incremental inductive learning, which they generally did (see Fig. 3A, Group 2). Thus, a significant part of Group 2’s attention may have remained on deductive reasoning. At the same time, the specific 1-5 cancer inducers (e.g., Acoseroc) were in effect a confound of the more general inductive understanding (the suffix only), such that if one only noticed the specifics, they could proceed with no obvious inconsistencies throughout learning and testing. Indeed, levels of confidence for the incremental learning (both inductive and deductive) and incremental deductive inference answers were highly similar (Fig. 3). Thus, Group 2 may have been led astray, so to speak, by experiencing the deductive component first, without recognizing the inductive pattern (suffix similarity).

At the same time, inductive performance was ultimately comparable between Groups 1 and 2. This may be due to the high degree of difficulty of the inductive component, with Group 1, who experienced the inductive component first, being more aware of the difficulty (leading to lowered confidence) without being able to do anything about it. In any case, our results show that actual understanding and one’s awareness of it indeed can diverge, which may arise from inherent confounds existing in complex, more natural scenarios.

### One-shot experience

For all participants combined, performance and confidence with the one-shot learning example (for both learning and inference) were relatively low (again, especially for inductive inference), and we found no effects of group (i.e., experience order) on either learning or inference. Thus, as might be expected, without proper rule extraction prior to the one-shot novel case, successful transfer could not occur.

For those who were able to extract the rule from the incremental experience (i.e., rule-catch group), they generally had success not only with the incremental cases, but also with the novel one-shot case (as reflected in the original ANOVA comparing rule-catch and not, the first regression showing the significant relation between rule and score, and in the breakdowns for one-shot inference, i.e., Fig. 3E&F). In fact, with the rule-catch group, we did find some evidence that the Group 1 order — inductive-then-deductive experience — helped to promote transfer of the rule to the one-shot problem. This order of experiencing the inductive component first may have helped to focus attention on the most challenging aspect of the problem, promoting the extraction of the suffix compound (e.g., ‘–seroc’) as the actual causal factor more strongly, enabling better transfer in the one-shot example. Kumaran ^24^ also found that stronger initial schema formation enabled more successful transfer to novel cases. Our findings are consistent with this, and thus suggest the extension to one-shot transfer, although future research is needed to verify possible facilitation from experiencing the inductive component first.

In any case, those who extracted the rule successfully during the incremental experience generally transferred it to the one-shot case — i.e., they also targeted the new suffix as the critical causal factor inducing the tumor, which enabled generalization to novel substances (i.e., ‘-cemel’). We thus found evidence for successful one-shot learning and generalization in a challenging logical reasoning paradigm, in which participants transferred their knowledge to the novel case even under the sparest of conditions: one instance of a novel case, i.e., one data point.

At the same time, however, their confidence lagged behind the performance scores. Thus, even though they transferred properly, they nonetheless were uncertain about it. Again, then, we found a decoupling between performance and confidence, but in the opposite direction of the prior case, in which actual knowledge here is accurate, but uncertainty about it remains high. Two general possibilities might explain the current mismatch. First, it is possible that the metacognitive system itself simply performs poorly: e.g., in the uncertainty derivations, the comparison among them, or control strength necessary to influence cognitive processing and behavior. The alternative possibility, however, is that the results actually reveal a well-designed cognitive system. That is, the results are consistent with an overall system that is operating with a set of hypotheses weighted by their individual certainties; and when required to select one of them, the leading hypothesis is chosen, even with low certainty^14,26^. Such a process is similar to ‘jumping to conclusions’ (i.e., revealing an overgeneralization bias)^26,27,28^, yet is critically different to the extent ‘jumping to conclusions’ is doing so without the corresponding caution (reflected in lower confidence) to prime updating of this knowledge in the future. Whereas the former may be seen as a clinical dysfunction, or an inherent flaw in the cognitive system^29^, the latter may reflect an optimized strategy: for example, in an environment where exploitation prospects are low (e.g., limited resources, high competition), random-exploration and trial-and-error learning too costly (e.g., heightened punishment due to scarce resources, competition, predation), with new opportunities that therefore must be seized upon when they present themselves, however minimal the evidence of how to exploit them. Evolutionary theorists have argued that this indeed was the type of environment that primates more generally, and humans in particular, evolved in^20,30,31^, although it also may reflect reinforcement contingencies for many people in our complex modern world. Computational analysis will be needed to help specify the conditions under which an apparent action bias toward generalization with minimal evidence, coupled with heightened metacognitive sensitivity, may provide a best-of-both-worlds cognitive strategy.

## Conclusions

When problems are relatively manageable, learning rate manipulation (by the cognitive control process) can be sufficient; but as problem difficulty increases, more elaborate means are needed for effective learning. We found that concentrated sequential experience (inductive-then-deductive or vice versa) proved to be far superior to intermixed (i.e., random) experience, and we also found some evidence that for those who were able to extract the rule (i.e., rule-catch group), inductive-then-deductive may have helped transfer the rule to the one-shot novel case, perhaps concentrating attention on the most difficult, inductive inference. As expected, we also found that those who extracted the rule properly, performed better and reported higher confidence than those who did not. At the same time, we also found uncertainty to be notably decoupled from actual knowledge. During sequential experience, those who experienced the deductive-then-inductive order were *overconfident*, which may have reflected a predominant focus on deductive reasoning, with a concomitant lack of awareness of the inductive component. In addition, for one-shot generalization, even though the rule-catch group showed superior ability to transfer their knowledge to the novel case, their confidence lagged. This finding appears to reflect a willingness to generalize with minimal experience, while maintaining skepticism about it — an overall strategy that may be particularly successful in difficult conditions that nonetheless harbor hidden opportunities.

Future research will be needed to further delineate and explain this complex relationship between causal reasoning and the metacognitive awareness of it. Indeed, our findings in turn suggest the need for explicit *manipulation* of one’s environmental experience to achieve a sufficient degree of controlled systematic experience with particularly challenging problems. Moreover, without someone to guide this (e.g., teacher), one normally must do it him/herself, in turn suggesting that an internal control process itself must orchestrate this manipulation of one’s own environmental experience^20,30^. Future computational development, together with behavioral and neurobiological studies are warranted to elucidate the possible more elaborate control mechanisms. Improvements to the current experimental design include a more explicit manipulation of problem difficulty, and confidence augmented by additional measures, such as response time (RT)^32^. Future work may also include the development of computational systems that can help overcome our blind spots and augment our ability to identify causal relationships, especially in particularly challenging problem domains^21^.

## Materials and Methods Participants

Seventy-seven adults (ages 20-33 years, mean ± standard deviation: 24 ± 2.5; 43 females) participated in the experiment. Participants with no reported history of neurological or psychological disorders were recruited from the local community around the Daejeon area via online job sites and college websites. With consideration of sex ratio, participants were divided into three groups based on learning order (described further below). Group 1, 2, and 3 consisted of twenty-six participants (ages 21-33, 24.42 ± 3.00; 15 females), twenty-six participants (ages 21-31, 24.08 ± 2.53; 14 females), and twenty-five participants (ages 20-28, ± 2.09; 14 females), respectively. All participants gave written consent, and the experimental procedures were approved by the Institutional Review Board (IRB) of Korea Advanced Institute of Science and Technology (KAIST).

### Sentence inference task

#### Background

*Inference* refers to the process of deriving a new proposition or judgment from one or more propositions or judgments that are known *a priori*. Typically, inference is divided into two types: inductive and deductive.

*Inductive inference* is a reasoning method that begins with collecting individual evidences and gradually finding general principles that can explain the individual cases. A famous example of inductive inference is as follows:

> *Premises: A. Socrates is mortal, B. Aristotle is mortal, C. Einstein is mortal. Conclusion: All men are mortal.*
>
> *Deductive inference* is a reasoning method to derive a logically certain conclusion from one or more propositions. An example of deductive inference is the law of syllogism, which is further subdivided into Modus ponens (law of detachment), Modus tollens (law of contrapositive), etc. A simple example of syllogism is as follows:
>
> *Premises: 1. All men are mortal, 2. Socrates is a man.*
>
> *Conclusion: 3. Therefore, Socrates is mortal.*

#### Sentence structure

To build a sentence structure that requires inference for acquiring information, we used schematized forms of inductive and deductive inference shown in Fig. 1. We generated the concepts, words and sentence structure with reference to the Korean translation of Principles of Cancer Biology^33^.

To encourage inductive inference, we made artificial words by using a common suffix. For example, substances that cause ‘Karmeictumor’ contain ‘–seroc’ in their name, such as Acoseroc, Benzoseroc, Uvoseroc, etc. In the same way, words whose suffix is ‘-myce’ cause ‘Parpenicumor’.

Deductive sentences were constructed by using a syllogism. For example, from the following three sentence sequences, we can infer the last one: ‘Humoseroc in rotten water induces Karmeictumor.’, ‘Karmeictumor is known as a tumor that triggers neck-ache.’, and ‘The tumor, which triggers neck-ache, easily induces metastasis and reinvasion.’ so that ‘Humoseroc can induce the tumor that easily induces metastasis.’. The sentence stimuli are shown in Table 1, although all were presented in Korean.

### Task paradigm

Every participant received incremental learning (both inductive and deductive parts), as well as one-shot learning, and all groups received incremental learning first, followed by one-shot learning. To test both type and order of experience, we divided participants into three groups: Group 1 received the inductive learning sentences first, followed by the deductive ones; Group 2 received vice versa (deductive prior to inductive); and Group 3 received all incremental learning sentences in Table 1 (i.e., deductive and inductive) in random order, and then all one-shot learning sentences in randomized order — and thus inductive and deductive learning being intermixed.

The task consisted of learning, exercise and test phases. In the learning phase, to promote incremental learning, the sentences denoted in Table 1 were presented 10 times each, with one caveat. For the inductive sentences, we created two versions of each sentence, one with “induced”, and the other with “did not induce”, with both sentence types also including the premise “In this case”: for example, “In this case, Acoseroc in tobacco smoke induced Karmeictumor by DNA impair” and “In this case, Acoseroc in tobacco smoke did not induce Karmeictumor by DNA impair”. The number of presentations of each sentence type was 8 (for “induced”) and 2 (for “did not induce”). This manipulation accomplished two objectives: (1) making the task sufficiently difficult to test our hypotheses with respect to more challenging logical reasoning problems; and (2) promoting uncertainty, one of the study’s central examination objectives. Thus, the total number of sentence stimuli was 143: 140 for Incremental Learning and 3 for One-shot Learning.

After the learning phase, an exercise phase was instantiated whereby the participants were asked two simple questions regarding their gender (“Are you a male?”) and major (“Does your major relate to biology or pharmacy?”) to (a) distract them to minimize contributions of short-term memory, and (b) provide familiarity with the question procedure. The test phase then consisted of thirty-five yes/no questions: those based on the exact information in the presented sentences (e.g., “Among the ingredients that cause Karmeictumor, Acoseroc is also included.”), called “Learning” questions, and new ones (e.g., “Nagoseroc will induce Karmeictumor.”), called “Inference” questions, which were not presented in the learning phase. The thirty-five test questions were of eight types (with the number of each question type in parenthesis): Incremental Inductive Learning (6), Incremental Inductive Inference (6), Incremental Deductive Learning (7), Incremental Deductive Inference (6), One-shot Inductive Learning (1), One-shot Inductive Inference (4), One-shot Deductive Inference (1), and One-shot Inductive + Deductive Inference (4), with an example of the last type being “Among the substances that cause intraepithelial carcinoma, there may be Yamincemel.” (see Fig. 1). The “Learning” questions used in the test portion were randomly selected to diminish the effect of familiarity for any particular sentence.

Finally, to rule out participants having superior memory capacity enabling them to potentially memorize all sentences without ever developing an inductive or deductive rule, we also tested them on a modified version of the 3-digit number span subtest from the WAIS (Wechsler Adult Intelligence Scale)^34^.

### Task procedure

First, participants were instructed as follows:

“In the first phase, sentences will be shown on the computer monitor for about 25 minutes. Imagine that you performed some kind of experiment and the shown sentences are the results of it. You need to learn it, and do not think about your previous knowledge. They will be repeatedly shown but please pay attention until it is over.

When the first phase is finished, you will receive a training phase with two questions. First, you can answer the questions by pressing O (for correct) or X (for incorrect). After answering each question, you will be asked to score how confident you are about your answer on a scale from −5 to +5. Move the point on the screen with ‘←’ or ‘→’ key and press Enter.

For the third phase, you will be asked a series of questions about what you learned in the first phase. The method of submitting your answers is the same as the second (training) phase. Some words that were not shown in the learning phase may appear, but you can infer an answer from what you learned in the first (learning) phase.”

The learning phase then initiated. Six to eight seconds were given randomly for reading each sentence and also one to four seconds randomly for the fixation period between sentences. After the learning phase, the participants were given the two exercise questions (gender and major) and then the thirty-five test questions. As shown in the instructions to the participants, they were asked to read the question sentence on the screen and answer with O/X for whether that sentence was correct or not. The computer screen then displayed the additional question asking ‘How confident are you about your answer?’ Participants then rated their confidence from −5 to 5. The exercise and test phase were self-paced.

When the main task was completed, a brief interview was conducted to determine whether the participant developed the inference rule or not. For example, we asked:

> *“How did you remember the causal relationship between the words?”*
>
> *“How did you answer the question that includes a new word you did not see in the first phase?”*
>
> *“Did you find some kind of rule?”* … etc.

After the interview, forty participants were further tested on the modified version of the WAIS memory test, with five trials performed by each participant, with each trial consisting of a serial presentation of 4 3-digit numbers to be memorized. Each 3-digit number appeared for two and a half seconds, with a half-second fixation period in between each number. Serial presentation of the four numbers was followed by a recall question to test whether they remembered each digit (ones, tens, or hundreds) of each number. Each question was shown for five seconds.

### Statistical analysis

Behavioral data were analyzed using IBM SPSS statistics 22.0 software. The average score and confidence rating (‘-5 to +5’ scale converted to ‘0-100’ via ‘50+10*rating’) of the three groups were compared using analysis of variance (ANOVA). When comparing two groups directly we used Student’s t-test. Finally, regression was conducted to quantify the correlation between variables.

## Acknowledgements

We greatly appreciate Dr. Jee Hang Lee’s assistance and insightful comments. This work was supported by Samsung Research Funding Center of Samsung Electronics under Project Number SRFC-TC1603-06, the research fund of the KAIST (Korea Advanced Institute of Science and Technology) (Grant code: G04150045), and the National Research Foundation of Korea(NRF) grant funded by the Korea government(MSIT) (No. 2017R1C1B2008972).

